# Intake of dietary fiber, fruits, and vegetables, and risk of diverticulitis

**DOI:** 10.1101/487462

**Authors:** Wenjie Ma, Long H. Nguyen, Mingyang Song, Manol Jovani, Po-Hong Liu, Yin Cao, Idy Tam, Kana Wu, Edward L. Giovannucci, Lisa L. Strate, Andrew T. Chan

**Author notes:** The authors contributed equally (Lisa L. Strate, Andrew T. Chan). **Corresponding author:** Dr. Andrew T. Chan, Clinical and Translational Epidemiology Unit, Massachusetts General Hospital, 55 Fruit Street, Boston, MA 02114.

## Abstract

**Background & Aims:** Although low fiber intake has been considered a risk factor for diverticulitis, prospective evidence is limited and conflicting, with little known about variation in the protective effects according to food sources. We assessed the associations of intakes of fiber and major food sources of fiber including fruits and vegetables with risk of diverticulitis.

**Methods:** We followed 50,019 women in the Nurses’ Health Study (1990-2014) and 48,292 men in the Health Professionals Follow-up Study (1986-2014) who were free of diverticulitis, cancer, and inflammatory bowel disease at baseline. Incident diverticulitis was identified through self-report with validity confirmed by review of medical records.

**Results:** During a mean follow-up time of 22 years, we documented 4,343 incident cases of diverticulitis in women and 1,142 cases in men. Compared to participants in the lowest quintile, the multivariable HRs (95% CIs) of diverticulitis in the highest quintile of total fiber intake were 0.86 (0.78-0.95; *P*-trend=0.002) among women and 0.63 (0.51-0.79; *P*-trend<0.001) among men. Fiber from different food sources, except for vegetable fiber in women, was associated with a decreased risk of diverticulitis. Furthermore, total whole fruit intake was associated with reduced risk of diverticulitis in both cohorts with a multivariable HR for diverticulitis of 0.95 (0.92-0.98; *P*-trend<0.001) in women and 0.91 (0.86-0.96; *P*-trend<0.001) in men for every serving increase of total whole fruit intake per day.

**Conclusions:** Higher intake of dietary fiber and fiber from different food sources are associated with a lower risk of diverticulitis. A greater intake of whole fruit is also associated with reduced risk.

## INTRODUCTION

Diverticulosis, or the presence of colonic diverticula, is very common in Western countries with prevalence increasing with age, reaching 60% by age 70.^1^ The principle complication of diverticulosis is diverticulitis, acute inflammation of diverticula and the surrounding colon, which can lead to significant complications that necessitate colectomy. Diverticulitis is among the leading gastrointestinal indications for hospitalizations and outpatient clinic visits in the United States.^2, 3^ Growing evidence has linked lifestyle and dietary factors to the etiopathogenesis of diverticulitis.^4, 5^

Fiber intake is perhaps the most studied dietary risk factor for diverticular disease, a collective term for both clinically significant and symptomatic diverticulosis and diverticulitis. Wide geographical and temporal differences in the prevalence of diverticular disease have been observed, and are thought to be secondary to differential intake of dietary fiber.^6^ Findings from our group and others have supported this hypothesis through prospective cohort studies.^5, 7–9^ However, colonoscopy-based studies have cast doubt on the relationship between a low-fiber diet and diverticulosis.^1, 10^ Furthermore, data on whether the protective effects of dietary fiber vary according to fiber subtypes and food sources are still limited and conflicting.^7–9^ Our earlier study in men found that the inverse association between fiber and symptomatic diverticular disease was primarily due to fruit and vegetable fiber^7^, whereas another UK study in women indicated that cereal and fruit fiber but not vegetable fiber were associated with reduced risk of diverticular disease^9^. Additionally, as major sources of dietary fiber, individual fruits and vegetables vary in fiber content and composition of other nutrients that may impact their beneficial effects. A recent genome-wide association analysis showed that genes associated with diverticular disease shared a common etiology with fresh fruit intake^11^, further suggesting a causal link. However, no prior study has examined what types of intake of fruits and vegetables may influence risk of diverticulitis. Therefore, we conducted a comprehensive, prospective evaluation of dietary fiber and fruit and vegetable intake in relation to risk of diverticulitis in two large cohorts, the Nurses’ Health Study (NHS) and Health Professionals Follow-up Study (HPFS).

## METHODS

### Study Population

The NHS is a cohort of 121,700 US female registered nurses aged 30 to 55 years at enrollment in 1976.^12^ The HPFS is a parallel cohort of 51,529 male health professionals aged 40 to 75 years at enrollment in 1986.^13^ Participants have been mailed questionnaires every two years since inception querying demographics, lifestyle factors, medical history, and disease outcomes, with a follow-up rate greater than 90% of available person-time. The study was approved by Institutional Review Boards of Brigham and Women’s Hospital and Harvard T.H. Chan School of Public Health. Return of the questionnaires was considered to imply written informed consent.

We excluded participants who reported a diagnosis of diverticulitis, non-melanoma cancers, or inflammatory bowel disease prior to baseline (1990 for the NHS and 1986 for the HPFS), those who had incomplete information for dietary data, and those who reported implausible total energy intake (< 500 or > 3500 kcal/day for the NHS and < 800 or > 4200 kcal/day for the HPFS). After exclusions, a total of 50,019 women in the NHS and 48,292 men in the HPFS were included in the primary analysis (**Supplemental Figure 1**).

### Dietary Assessment

Dietary intake data were assessed through administration of a validated 131-item semi-quantitative food frequency questionnaire (FFQ) every 4 years, in which participants were asked how often they typically consumed each food of a standard portion size during the previous year. We included all whole fruits and all vegetables on the FFQ in our analyses, and fruit juice including apple, orange, grapefruit, and other juice was considered separately as it typically includes added sugar and is associated with an increased risk of adverse health outcomes such as diabetes^14^ and greater weight gain^15^. Fruits and vegetables with similar culinary usages and nutrient profiles were combined, e.g., apples and pears. We ranked fruits and vegetables by their proportional contributions to total intake of fruit fiber or vegetable fiber based on results from all available FFQs in the NHS and HPFS.

Daily intake for each nutrient was calculated by multiplying the reported frequency of each food item by its nutrient content, and then summing across foods. Fiber intake was calculated using the Association of Official Analytical Chemists method (accepted by the US Food and Drug Administration and the Food and Agriculture Organization of the World Health Organization for nutrition labeling purposes).^16^ Data on soluble fiber and insoluble fiber were available directly from some manufactures, calculations from ingredients (personal communication from USDA), analysis done by Kellogg’s specifically from the Channing laboratory, as well as other dietary fiber food composition data (Horvath and Robertson 1986, USDA 1993).^8^ We adjusted fiber intake for total caloric intake using the nutrient residual method.^17^ Fiber intake from major food sources, including cereals, vegetables, and fruits, was also considered separately. FFQs have shown good reproducibility and validity for assessing intake of fiber, fruits, and vegetables.^18, 19^ With a correction for the ratio of the within-person to the between-person variation, the correlation coefficient comparing diet assessment from FFQ with multiple 7-day dietary records was 0.68 for dietary fiber and ranged from 0.38 (strawberries) to 0.95 (bananas) for individual fruits (median: 0.76) and 0.25 (kale, mustard greens or chard greens) to 0.73 (lettuce) for individual vegetables (median: 0.46).

### Ascertainment of Diverticulitis

In the NHS, participants were asked in 2008 and 2012 if they ever had a diagnosis of diverticulitis requiring antibiotic therapy or hospitalization. If yes, participants were subsequently asked the year of each episode dating back to 1990. In 2014, participants were asked the same question, but restricted to the last two years. In a review of 107 medical records from women reporting incident diverticulitis on the 2008 questionnaire, self-report was confirmed in 88% of cases. In the HPFS, beginning in 1990, participants who reported newly diagnosed diverticulitis or diverticulosis on the biennial study questionnaires were sent supplementary questionnaires that ascertained the date of diagnosis, presenting symptoms, diagnostic procedures, and treatment for each reported event. Diverticulitis was defined as abdominal pain attributed to diverticular disease and one of the following criteria: diverticular complications including perforation, abscess, fistula, or obstruction; hospitalization, antibiotic therapy, or surgery resulting from diverticulitis; pain categorized as severe or acute; or abdominal pain presenting with fever, requiring medical therapy or radiologic evaluation with an abdominal computed tomography. The validity of self-reported diverticulitis in HPFS was confirmed previously^20, 21^ with 84% of diverticulitis cases confirmed by chart review. Beginning in 2006, we revised our supplementary questionnaire to further assess uncomplicated diverticulitis, diverticular complications including abscesses, fistula formation, perforation, obstruction, diverticular bleeding, and asymptomatic diverticulosis.

### Assessment of Covariates

Participants reported height at the time of enrollment. At baseline and updated biennially, information on body weight, smoking status, menopausal status and menopausal hormone use (women only), physical examination, and use of multivitamins, aspirin, other nonsteroidal anti-inflammatory drugs (NSAIDs), or acetaminophen were obtained. Body mass index (BMI) was calculated as weight in kilograms divided by height in meters squared. Physical activity was assessed every 2-4 years using validated questionnaires.^22^ Hypertension and hypercholesterolemia were self-reported, with their validity previously confirmed.^23^ In our analysis, we allowed covariates to be time-varying using the most recent information.

### Statistical Analysis

Person-time was calculated from the date of baseline questionnaire until the date of diagnosis of diverticulitis, death, last follow-up questionnaire, or the end of the study period (June 2014 for NHS or January 2014 for HPFS), whichever came first. We censored participants who reported a new diagnosis of gastrointestinal cancer or inflammatory bowel disease at the date of diagnosis. We used simple updating or the most recent dietary information available (i.e., the intake reported on the most recent FFQ at the start of each 2-year follow-up interval). Missing values for a given FFQ were carried forward from prior available assessments.

Participants were categorized into quintiles according to their energy-adjusted fiber intake. Linear trend was assessed by assigning the median value to each category and modeling this as a continuous variable. Using Cox proportional hazards regression with time-varying dietary intake and covariates, we estimated hazard ratios (HRs) and 95% confidence intervals (CIs).

Proportional hazards assumption was evaluated by testing the significance of the interaction term between exposure and age by using the Wald test, and no violation of the proportional hazards assumption was observed (*P* for interaction > 0.05).

We stratified the analysis jointly by age at the start of follow-up and calendar time of the current questionnaire cycle to control for confounding by age, calendar time, and any possible two-way interactions between these two timescales. In multivariate analysis, we adjusted for BMI, menopausal status and menopausal hormone use (women only), vigorous physical activity (METs ≥ 6, including jogging, running, bicycling, swimming, tennis, squash or racquetball, rowing, and heavy outdoor work), alcohol intake, smoking, use of aspirin, other NSAIDs, or acetaminophen, multivitamin use, recent physical examination as a proxy for healthcare engagement, hypertension, hypercholesterolemia, calorie intake, and red meat intake. We further controlled for total whole fruit intake for our analysis of individual fruits and total vegetable intake when testing individual vegetables for any independent association.

Statistical analyses were conducted using SAS software version 9.4 (SAS Institute Inc., Cary, NC). All *p*-values are 2-sided and < 0.05 was considered statistically significant.

## RESULTS

At baseline, women in the NHS and men in the HPFS had a mean age of 55 and 52 years, respectively. Participant characteristics according to quintiles of energy-adjusted dietary fiber intake are shown in **Table 1**. The average total fiber intake was 18.0 g for women and 21.1 g for men. In both cohorts, total fiber intake was positively correlated with age, physical activity, multivitamin use, recent history of physical examination, hypercholesterolemia, and diabetes, and was inversely correlated with BMI, alcohol intake, acetaminophen use, other NSAID use, and current smoking. Participants who had higher fiber intake tended to consume more fruit, vegetables, and whole grains but less red meat, resulting in a higher healthy eating index score.

**Table 1.**
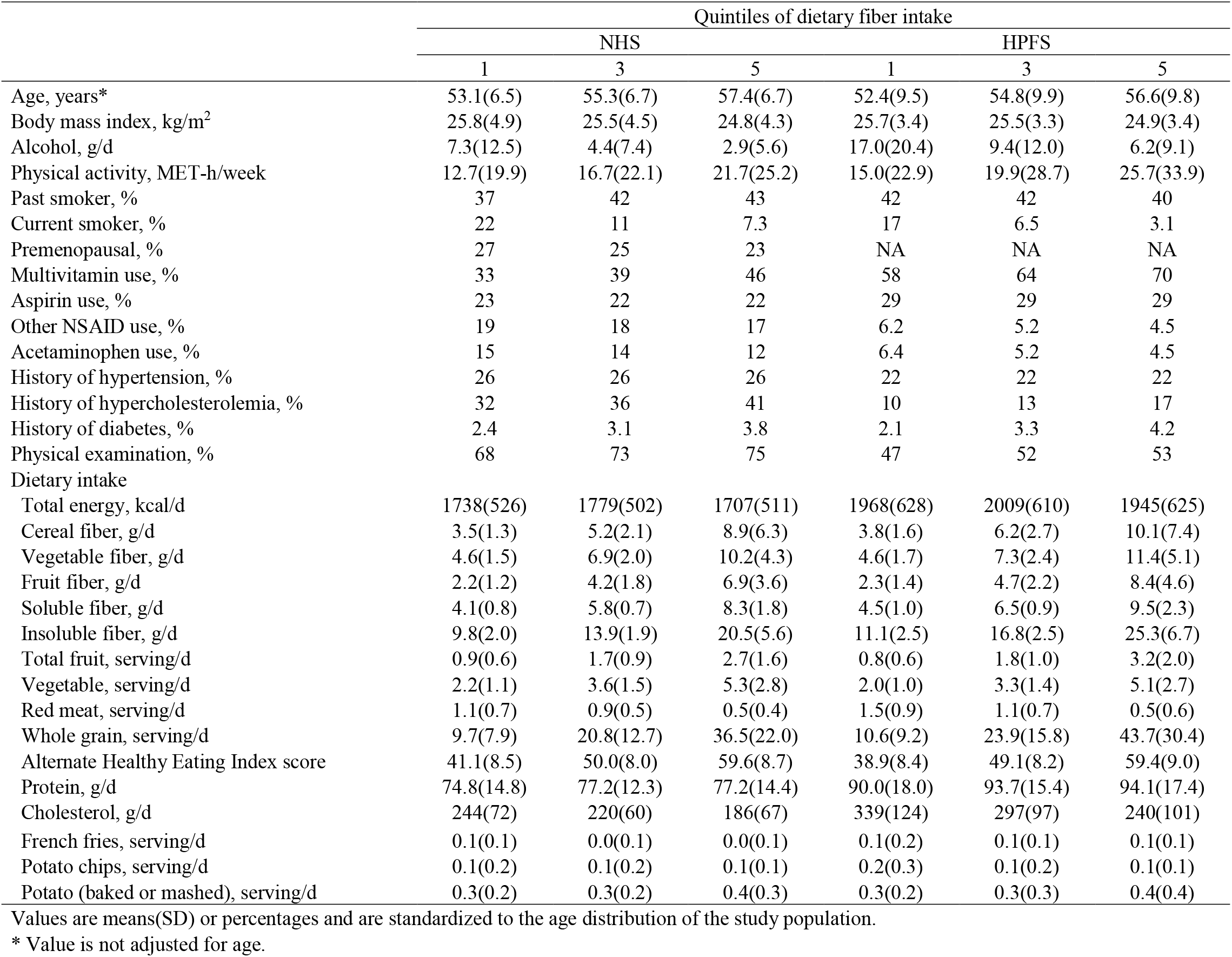
Baseline age-adjusted characteristics of participants according to energy-adjusted dietary fiber intake in Nurses’ Health Study (1990) and Health Professionals Follow-up Study (1986)

In the NHS, we documented a total of 4,343 incident cases of diverticulitis over 24 years, encompassing 1,106,402 person-years of follow-up. In age-adjusted analyses, higher intake of total fiber and fiber from cereals, fruit and vegetables were each associated with reduced risk of diverticulitis (**Table 2**). After further adjusting for other risk factors, including red meat intake, the associations between incident diverticulitis and total, cereal, and fruit fiber did not materially change, whereas the association with vegetable fiber was attenuated. Compared to women in the lowest quintile of dietary intake, the multivariable HRs (95% CIs) in the highest quintile were 0.86 (0.78-0.95; *P*-trend = 0.002) for total fiber, 0.90 (0.81-0.99; *P*-trend = 0.03) for cereal fiber, 0.83 (0.75-0.92; *P*-trend < 0.001) for fruit fiber, and 0.92 (0.83-1.01; *P*-trend = 0.31) for vegetable fiber.

**Table 2.** Energy-adjusted dietary fiber intake and risk of diverticulitis in Nurses’ Health Study (1990-2014)

|  | Quintiles of dietary intake |  |  |  |  | P for trend |
| --- | --- | --- | --- | --- | --- | --- |
|  | 1 | 2 | 3 | 4 | 5 |  |
| Total fiber |  |  |  |  |  |  |
| Median, g/d | 13.0 | 16.3 | 18.9 | 21.8 | 27.0 |  |
| No. of cases | 862 | 874 | 896 | 879 | 832 |  |
| Age | 1 (ref) | 0.96 (0.87, 1.05) | 0.94 (0.86, 1.04) | 0.90 (0.81, 0.98) | 0.82 (0.74, 0.90) | <0.001 |
| Multivariable* | 1 (ref) | 0.96 (0.88, 1.06) | 0.95 (0.87, 1.05) | 0.91 (0.83, 1.01) | 0.86 (0.78, 0.95) | 0.002 |
| Multivariable+red meat | 1 (ref) | 0.96 (0.88, 1.06) | 0.95 (0.86, 1.05) | 0.91 (0.83, 1.01) | 0.86 (0.78, 0.95) | 0.002 |
| Cereal fiber |  |  |  |  |  |  |
| Median, g/d | 2.9 | 4.3 | 5.6 | 7.0 | 9.8 |  |
| No. of cases | 813 | 880 | 895 | 903 | 852 |  |
| Age | 1 (ref) | 0.99 (0.90, 1.09) | 0.99 (0.90, 1.09) | 0.96 (0.87, 1.06) | 0.86 (0.78, 0.95) | 0.001 |
| Multivariable* | 1 (ref) | 0.98 (0.89, 1.08) | 0.99 (0.90, 1.09) | 0.98 (0.89, 1.08) | 0.90 (0.81, 0.99) | 0.02 |
| Multivariable+red meat | 1 (ref) | 0.98 (0.89, 1.08) | 0.99 (0.90, 1.10) | 0.98 (0.89, 1.08) | 0.90 (0.81, 0.99) | 0.03 |
| Fruit fiber |  |  |  |  |  |  |
| Median, g/d | 1.4 | 2.6 | 3.8 | 5.2 | 7.7 |  |
| No. of cases | 898 | 923 | 890 | 862 | 770 |  |
| Age | 1 (ref) | 0.99 (0.90, 1.08) | 0.97 (0.88, 1.06) | 0.89 (0.81, 0.98) | 0.79 (0.71, 0.87) | <0.001 |
| Multivariable* | 1 (ref) | 0.99 (0.90, 1.09) | 0.98 (0.89, 1.08) | 0.92 (0.84, 1.01) | 0.83 (0.75, 0.92) | <0.001 |
| Multivariable+red meat | 1 (ref) | 0.99 (0.90, 1.09) | 0.98 (0.89, 1.08) | 0.92 (0.83, 1.01) | 0.83 (0.75, 0.92) | <0.001 |
| Vegetable fiber |  |  |  |  |  |  |
| Median, g/d | 3.1 | 4.6 | 5.8 | 7.2 | 10.0 |  |
| No. of cases | 954 | 834 | 848 | 889 | 818 |  |
| Age | 1 (ref) | 0.93 (0.84, 1.02) | 0.94 (0.85, 1.03) | 1.01 (0.92, 1.10) | 0.90 (0.82, 0.99) | 0.14 |
| Multivariable* | 1 (ref) | 0.93 (0.85, 1.02) | 0.94 (0.86, 1.04) | 1.02 (0.93, 1.11) | 0.91 (0.83, 1.01) | 0.27 |
| Multivariable+red meat | 1 (ref) | 0.93 (0.85, 1.02) | 0.94 (0.86, 1.04) | 1.02 (0.93, 1.12) | 0.92 (0.83, 1.01) | 0.31 |

\* Further adjusted for body mass index, menopausal status and menopausal hormone use, vigorous activity, alcohol intake, smoking, aspirin use, other nonsteroidal anti-inflammatory drug use, multivitamin use, acetaminophen use, physical examination, hypertension, hypercholesterolemia, calorie intake, and red meat intake. Results for cereal, fruit, and vegetable fiber were similar if further mutually adjusting for fiber from other sources (i.e., except for the one under examination).

In the HPFS, we documented 1,142 incident cases over 28 years, encompassing 1,022,391 person-years of follow-up. Associations between fiber intake and incidence of diverticulitis in the HPFS were similar, but generally stronger compared to those in the NHS (**Table 3**). In age-adjusted analyses, higher intake of total fiber and fiber from cereal, fruit, and vegetables were each associated with reduced risk of diverticulitis. Multivariate adjustment attenuated, but did not materially change these observations. Compared to men in the lowest quintile, the corresponding HRs (95% CIs) in the highest quintile were 0.63 (0.51-0.79; *P*-trend < 0.001) for total fiber, 0.84 (0.69-1.02; *P*-trend = 0.03) for cereal fiber, 0.80 (0.65-0.98; *P*-trend = 0.02) for fruit fiber, and 0.73 (0.60-0.88; *P*-trend < 0.001) for vegetable fiber.

**Table 3.** Energy-adjusted dietary fiber intake and risk of diverticulitis in Health Professional Follow-Up Study (1986-2014)

|  | Quintiles of dietary intake |  |  |  |  | P for trend |
| --- | --- | --- | --- | --- | --- | --- |
|  | 1 | 2 | 3 | 4 | 5 |  |
| Total fiber |  |  |  |  |  |  |
| Median, g/d | 14.7 | 18.8 | 22.1 | 25.9 | 32.5 |  |
| No. of cases | 281 | 245 | 227 | 225 | 164 |  |
| Age | 1 (ref) | 0.84 (0.71, 1.00) | 0.76 (0.64, 0.91) | 0.75 (0.63, 0.89) | 0.55 (0.45, 0.66) | <0.001 |
| Multivariable* | 1 (ref) | 0.84 (0.70, 0.99) | 0.77 (0.64, 0.92) | 0.76 (0.63, 0.92) | 0.58 (0.47, 0.71) | <0.001 |
| Multivariable+red meat | 1 (ref) | 0.84 (0.71, 1.00) | 0.78 (0.65, 0.93) | 0.79 (0.65, 0.95) | 0.63 (0.51, 0.79) | <0.001 |
| Cereal fiber |  |  |  |  |  |  |
| Median, g/d | 3.1 | 4.7 | 6.3 | 8.1 | 11.4 |  |
| No. of cases | 245 | 238 | 261 | 193 | 205 |  |
| Age | 1 (ref) | 0.93 (0.78, 1.11) | 0.95 (0.80, 1.13) | 0.72 (0.60, 0.87) | 0.74 (0.62, 0.90) | <0.001 |
| Multivariable* | 1 (ref) | 0.93 (0.77, 1.11) | 0.95 (0.79, 1.14) | 0.73 (0.60, 0.89) | 0.78 (0.64, 0.95) | 0.002 |
| Multivariable+red meat | 1 (ref) | 0.93 (0.78, 1.12) | 0.97 (0.81, 1.16) | 0.76 (0.62, 0.92) | 0.84 (0.69, 1.02) | 0.03 |
| Fruit fiber |  |  |  |  |  |  |
| Median, g/d | 1.5 | 3.0 | 4.4 | 6.1 | 9.4 |  |
| No. of cases | 262 | 245 | 233 | 219 | 183 |  |
| Age | 1 (ref) | 0.95 (0.80, 1.13) | 0.87 (0.73, 1.04) | 0.80 (0.66, 0.95) | 0.68 (0.56, 0.82) | <0.001 |
| Multivariable* | 1 (ref) | 0.96 (0.81, 1.15) | 0.90 (0.75, 1.08) | 0.84 (0.69, 1.01) | 0.74 (0.60, 0.90) | <0.001 |
| Multivariable+red meat | 1 (ref) | 0.97 (0.81, 1.16) | 0.92 (0.76, 1.10) | 0.87 (0.72, 1.05) | 0.80 (0.65, 0.98) | 0.02 |
| Vegetable fiber |  |  |  |  |  |  |
| Median, g/d | 3.6 | 5.3 | 6.7 | 8.6 | 12.2 |  |
| No. of cases | 270 | 222 | 267 | 203 | 180 |  |
| Age | 1 (ref) | 0.86 (0.72, 1.02) | 1.00 (0.84, 1.18) | 0.75 (0.62, 0.90) | 0.66 (0.55, 0.80) | <0.001 |
| Multivariable* | 1 (ref) | 0.86 (0.72, 1.03) | 1.01 (0.85, 1.20) | 0.76 (0.64, 0.92) | 0.69 (0.57, 0.83) | <0.001 |
| Multivariable+red meat | 1 (ref) | 0.86 (0.72, 1.03) | 1.02 (0.86, 1.21) | 0.78 (0.65, 0.94) | 0.73 (0.60, 0.88) | <0.001 |

\* Further adjusted for body mass index, vigorous activity, alcohol intake, smoking, aspirin use, other nonsteroidal anti-inflammatory drug use, multivitamin use, acetaminophen use, physical examination, hypertension, hypercholesterolemia, calorie intake, and red meat intake. Results for cereal, fruit, and vegetable fiber were similar if further mutually adjusting for fiber from other sources (i.e., except for the one under examination).

Total whole fruit intake was associated with a reduced risk of diverticulitis in both cohorts (**Figure 1**). The multivariable HR (95% CI) of diverticulitis was 0.95 (0.92-0.98; *P* < 0.001) in the NHS and 0.91 (0.86-0.96; *P* < 0.001) in the HPFS for every serving increase of total whole fruit intake per day. When individual fruits were evaluated, increased intake of apples or pears consistently showed a significant or a trend towards significant association with lower risk of diverticulitis in both NHS (HR, 0.85; 95% CI, 0.76-0.96; *P* = 0.009) and HPFS (HR, 0.84; 95% CI, 0.69-1.03; *P* = 0.10), independent of total whole fruit intake. Inverse association was also observed between prunes and diverticulitis in the NHS (HR, 0.83; 95% CI, 0.69-0.99; *P* = 0.04), whereas none of the other fruits were significantly associated with diverticulitis in the HPFS. In contrast to the inverse association seen for whole fruit, fruit juice was not significantly associated with risk of diverticulitis, with a multivariable HR (95% CI) of 1.04 (0.99-1.08; P-trend = 0.10) in the NHS and 0.99 (0.91-1.07; P-trend = 0.73) in the HPFS for every serving increase of fruit juice per day.

**Figure 1.**
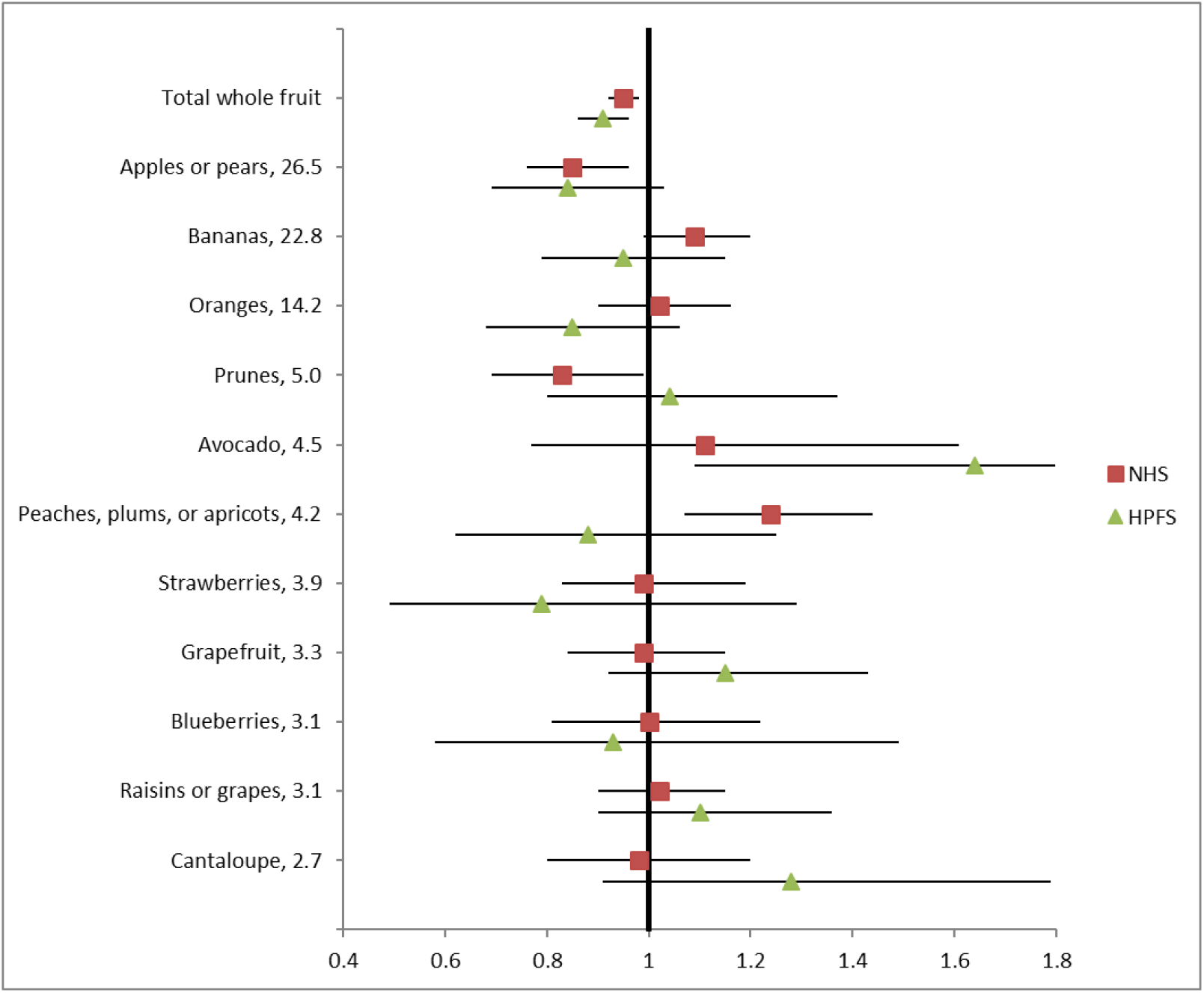
Total whole and individual fruit intake and risk of diverticulitis in NHS (1990-2014) and HPFS (1986-2014). The values following fruit names represented percentage of contribution to total fruit fiber based on pooled results from NHS and HPFS. Total whole fruit does not include fruit juice. Hazard ratios were shown for increase of one serving per day, adjusted for age, body mass index, menopausal status and menopausal hormone use (women only), vigorous activity, alcohol intake, smoking, aspirin use, other nonsteroidal anti-inflammatory drug use, multivitamin use, acetaminophen use, physical examination, hypertension, hypercholesterolemia, calorie intake, and red meat intake. Results for individual fruits were further adjusted for total whole fruit intake.

Increased total vegetable intake was associated with a reduced risk of diverticulitis in men (HR, 0.96; 95% CI, 0.92-0.99; *P* = 0.009), but not in women (HR, 0.99; 95% CI, 0.97-1.01; *P* = 0.17) (**Figure 2**). Among individual vegetables, mixed vegetable intake showed an inverse association with diverticulitis independent of total vegetable intake, in the HPFS, but not in the NHS, with HR (95% CI) of 0.55 (0.36-0.84; *P* = 0.006) for every serving increase of intake per day. We also observed a trend towards an inverse relationship between diverticulitis and beans or lentils in both women (HR, 0.78; 95% CI, 0.61-1.01; *P* = 0.06) and men (HR, 0.76; 95% CI, 0.50-1.14; *P* = 0.19).

**Figure 2.**
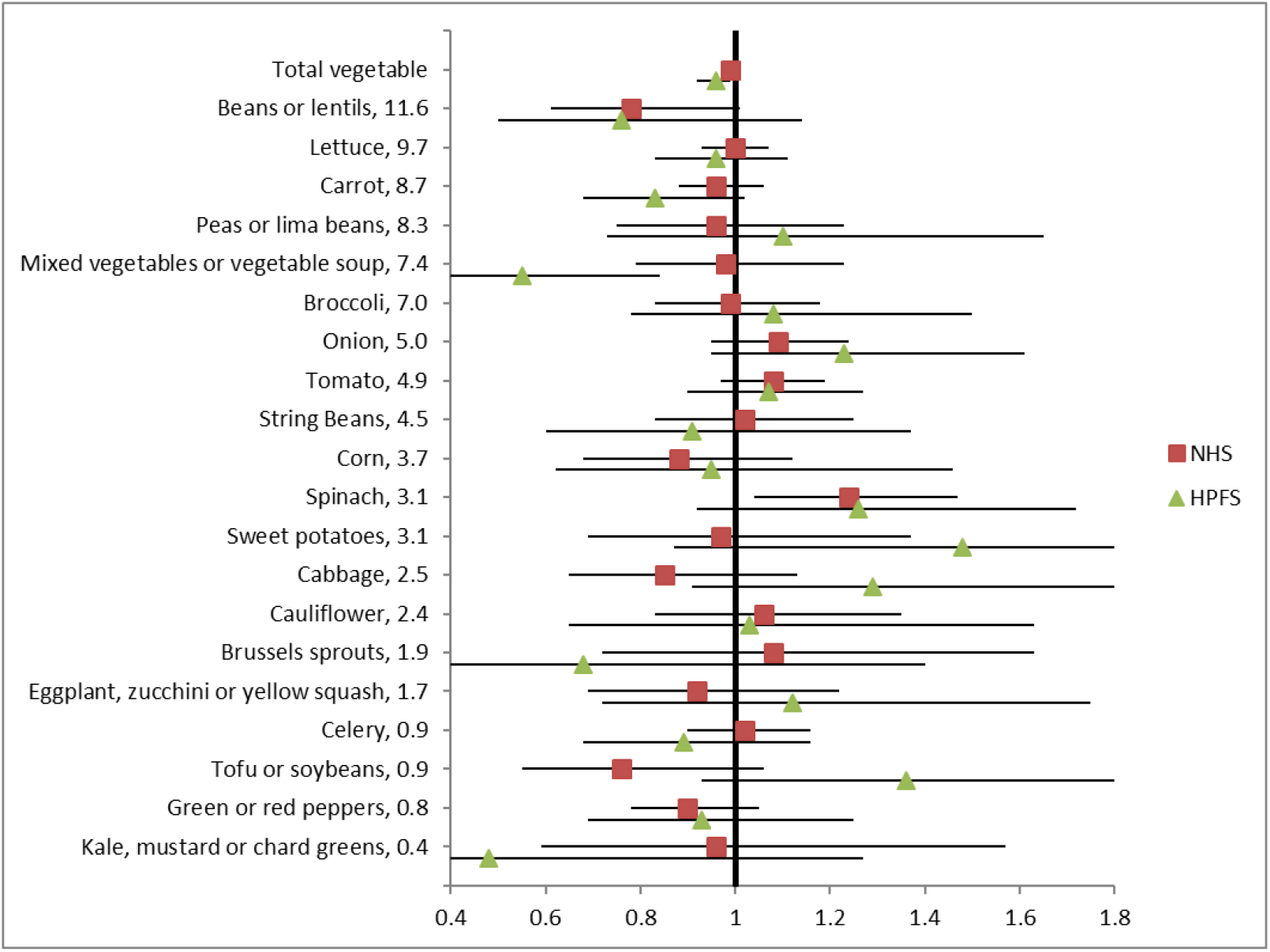
Total and individual vegetable intake and risk of diverticulitis in NHS (1990-2014) and HPFS (1986-2014). The values following vegetable names represented percentage of contribution to total vegetable fiber based on pooled results from NHS and HPFS. Hazard ratios were shown for increase of one serving per day, adjusted for age, body mass index, menopausal status and menopausal hormone use (women only), vigorous activity, alcohol intake, smoking, aspirin use, other nonsteroidal anti-inflammatory drug use, multivitamin use, acetaminophen use, physical examination, hypertension, hypercholesterolemia, calorie intake, and red meat intake. Results for individual vegetables were further adjusted for total vegetable intake.

Evaluating soluble and insoluble fiber separately, insoluble fiber showed a stronger association with risk of diverticulitis compared to soluble fiber in women, whereas the associations were similar in men (**Supplemental Table 1**). In the NHS, compared to women in the lowest quintile of dietary intake, the multivariable HRs (95% CIs) in the highest quintile were 0.95 (0.86-1.05; *P*-trend = 0.7 for soluble fiber and 0.86 (0.78-0.95; *P*-trend = 0.005) for insoluble fiber. The corresponding HRs (95% CIs) in the HPFS were 0.73 (0.59-0.89; *P*-trend = 0.001) and 0.77 (0.63-0.94; *P*-trend = 0.005), respectively.

We also separately examined the relationship between consumption of some other processed vegetable foods (that were not accounted into total vegetable intake), including potatoes (baked or mashed), potato chips, and French fries and diverticulitis. We found that after adjusting for other risk factors including fiber intake, intake of baked or mashed potatoes (HR for every serving increase of intake per day: 1.20; 95% CI: 1.07-1.36; *P* = 0.003), but not French fries (HR: 0.98; 95% CI: 0.63-1.51; *P* = 0.91) or potato chips (HR: 1.13; 95% CI: 0.96-1.32; *P* = 0.13) were associated with a modest increase in the risk of diverticulitis in the NHS, whereas none of these foods were significantly associated with diverticulitis in the HPFS.

## DISCUSSION

In two prospective cohorts of US women and men, higher fiber intake was associated with reduced risk of incident diverticulitis. Fiber from different food sources was also inversely associated with diverticulitis, with the exception of vegetable fiber in women. Additionally, greater consumption of total whole fruit and specific fruits such as apples or pears were consistently associated with a lower risk of diverticulitis.

The inverse association between fiber intake and diverticulitis is consistent with findings from previous prospective studies. In an earlier analysis utilizing only a subset of our current study population over relatively short-term follow-up (1988 to 1992), Aldoori et al reported that total fiber intake and fiber from fruit and vegetables, but not cereal, were associated with reduced risk of symptomatic diverticular disease in men.^7^ Using the EPIC-Oxford cohort, Crowe et al confirmed the inverse association between fiber intake and risk of diverticular disease requiring hospitalization or as the cause of death among men and women in the UK.^5^ It was subsequently demonstrated that a higher intake of dietary fiber, specifically fruit fiber and cereal fiber, was associated with a reduction in the risk of diverticular disease among UK women.^9^ However, a potential drawback with these studies is the inability to distinguish between diverticulitis, diverticular bleeding, and symptomatic uncomplicated diverticulosis; additionally, the two aforementioned UK studies were based on registry-level data and included only hospitalized patients; the influence of dietary fiber intake on less severe and more common presentations^24^ remains unclear. In contrast to findings from prospective studies of diverticulitis or diverticular disease requiring hospitalization, low intake of dietary fiber was not associated with increased risk of uncomplicated diverticulosis in cross-sectional, colonoscopy-based studies^1, 10, 25^. The divergent associations with diverticulitis and diverticulosis suggest that fiber may not be associated with the development of diverticulosis, but may play a role in preventing the inflammation associated with diverticulitis. Furthermore, data from the colonoscopy-based studies may not be comparable to our findings since diet was assessed after colonoscopy and patients may have modified their diet due to early symptoms or had differential recall according to the awareness of the disease. Our study extends previous literature by offering evidence that fiber intake is associated with reduced risk of diverticulitis in men and women across the spectrum of diverticulitis by leveraging prospective and repeated dietary assessments with long-term follow-up.

The associations of fruits and vegetables with diverticular disease have also been investigated in other observational studies, but results have been less consistent. In a case-control study conducted in Greece, patients with radiologically-confirmed diverticulosis had less frequent consumption of vegetables compared to controls, whereas the frequency of fruit consumption was not significantly different.^26^ Another case-control study in Taiwan, however, found consumption frequency of fruit and vegetables was not associated with right-sided asymptomatic diverticulosis.^27^ While these studies were limited by the retrospective design and small sample size, it is also likely that diverticulosis, in particular right-sided diverticulosis which is predominant in Asia, has different pathophysiology from left-sided diverticular disease or diverticulitis.^28^ An earlier analysis from the HPFS cohort demonstrated energy-adjusted associations of symptomatic diverticular disease with some fruits and vegetables, such as romaine or leafy lettuce, peaches, apricots, or plums, oranges, apples, and blueberries.^7^ Due to high correlations between individual and total fruits or vegetables, it is critical to control for total fruit or vegetable intake in the evaluation of any independent relationship between individual fruits or vegetables and diverticulitis. Our findings indicate that overall intake of fruit or vegetables may be more important compared to individual items in influencing the risk of diverticulitis. Apples or pears did appear to have the strongest individual associations, but this may reflect that these items were by far the most commonly consumed, contributing to 26.5% of total fruit fiber. Meanwhile, differences in contributions to fiber intake did not appear to account for the association of specific fruits and vegetables with disease risk.

Sex differences of the associations of fiber subtypes with diverticulitis were also consistent with the literature.^7–9^ In women, in parallel with the more pronounced association of fruit fiber and to a lesser extent, cereal fiber, with risk of diverticulitis compared to vegetable fiber, we also observed a stronger association for insoluble vs. soluble fiber. Insoluble fiber, the predominant fraction of dietary fiber, is most abundant in whole grain foods and could help prevent constipation. In men, however, the association of dietary fiber and diverticulitis was primarily due to fruit and vegetable fiber. Overall, fruit and vegetables tend to be higher than cereals in cellulose^29^, the fiber subtype most strongly associated with diverticular disease in prior studies^8^. This might partially explain the beneficial effects of a higher fiber intake from fruit and vegetables and a higher intake of fruit and vegetables we observed in men. Yet, these results should be carefully interpreted as the distribution of total fiber between soluble and insoluble subtypes is dependent on the method of analysis.^30^

The biological mechanism through which dietary fiber may decrease risk of diverticular disease was initially hypothesized to be mediated by its effects on colonic motility and intraluminal pressure.^31, 32^ More recent evidence suggests that fiber alleviates obesity-induced chronic inflammation and also has weight-unrelated anti-inflammatory effects, the latter of which is likely mediated through intestinal microbiota.^33, 34^ Meanwhile, sex hormones have been shown to alter intestinal permeability^35^ and the gut microbiome^36^. A prior study suggested that menopausal hormone therapy was associated with increased risk of diverticular disease in women.^5^ Studies also showed that of the three sources of fiber, fruit fiber had the strongest inverse association with circulating concentrations of estradiol, followed by grain fiber; in contrast, vegetable fiber was not associated.^37^ Taken together, fiber from different food sources may have divergent interactions with sex hormones and gut microbiome that in turn differentially influence risk of diverticulitis in women compared with men. However, future investigations are warranted to verify our hypothesis and better understand the potentially varying roles of fiber subtypes and food sources in the development of the disease.

The strengths of our study include the prospective design, detailed and repeated collection of diet and lifestyle information, long-term follow-up and ascertainment of diverticulitis managed in the inpatient and outpatient setting. Moreover, detailed, contemporaneous information on potential confounders was collected in parallel with fiber, fruit, and vegetable intake.

Several limitations are worth noting. First, the ascertainment of dietary intake and diverticulitis were based on self-report. However, the validity of our dietary instruments has previously been demonstrated^18, 19^, and confirmation of self-reported diverticulitis is high. Second, the diagnosis of diverticulitis in the NHS was based on recall among participants who responded to the 2008, 2012, and 2014 questionnaires, which we believe will not result in bias because our outcome is unlikely to be fatal.^38^ Finally, our cohorts included mostly Caucasian health professionals, and future studies in other ethnic groups are needed considering potential ethnic disparities in diverticular disease.

In conclusion, a higher intake of fiber is associated with a lower incidence of diverticulitis. Potential sex differences exist in the effects of fiber from different food sources. Total whole fruit intake and, to a lesser extent, vegetable intake, is also associated with a reduced risk. Our findings lend support for public health recommendations of maintaining sufficient fiber intake from diet and provide practical dietary guidance for patients in the prevention of diverticulitis.

## Supporting information

## Author contributions

Study concept and design: WM, LLS, ATC; acquisition of data: KW, ELG, LLS, ATC; statistical analysis: WM; interpretation of data: all authors; drafting of the manuscript: WM, LLS, ATC; critical revision of the manuscript for important intellectual content: all authors.

## Acknowledgements

We would like to thank the participants and staff of the Nurses’ Health Study and Health Professionals Follow-up Study for their valuable contributions.

