## Supplementary material for "Intake of dietary fiber, fruits, and vegetables, and risk of diverticulitis"

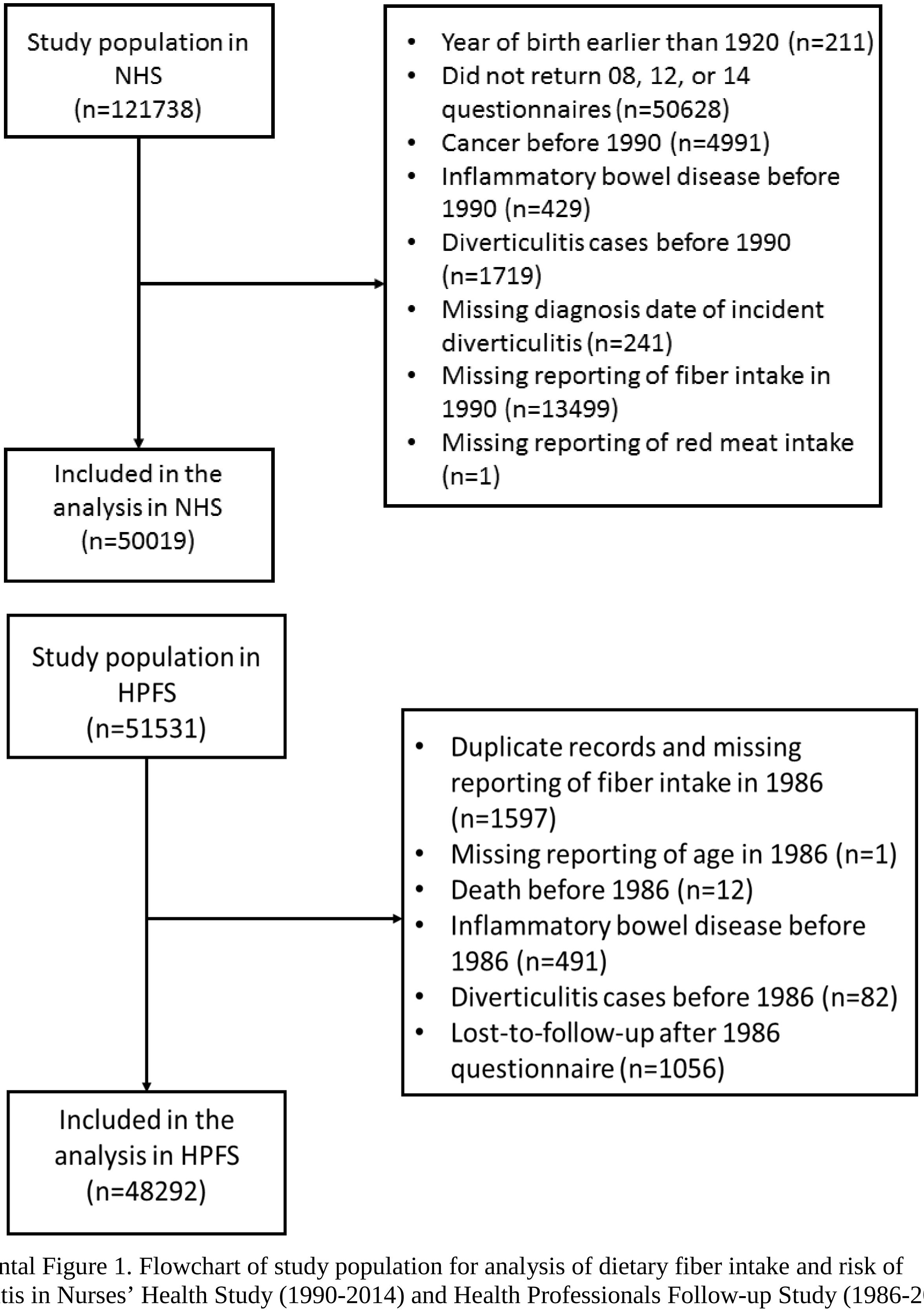


Supplemental Figure 1. Flowchart of study population for analysis of dietary fiber intake and risk of diverticulitis in Nurses' Health Study (1990-2014) and Health Professionals Follow-up Study (1986-2014)
