## Supplementary material for "Intake of dietary fiber, fruits, and vegetables, and risk of diverticulitis"

Supplemental Table 1. Energy-adjusted soluble and insoluble fiber intake and risk of diverticulitis in Nurses' Health Study (1990-2014) and Health Professional Follow-Up Study (1986-2014)

| | Quintiles of dietary intake | | | | | |
|---|---|---|---|---|---|---|
| | 1 | 2 | 3 | 4 | 5 | P for trend |
| Soluble fiber | | | | | | |
| NHS | | | | | | |
| Median, g/d | 3.7 | 4.6 | 5.3 | 6.2 | 7.9 | |
| No. of cases | 918 | 778 | 868 | 885 | 894 | |
| Age | 1 (ref) | 0.94 (0.86, 1.04) | 0.93 (0.84, 1.02) | 0.96 (0.88, 1.06) | 0.92 (0.84, 1.01) | 0.17 |
| Multivariable* | 1 (ref) | 0.95 (0.86, 1.04) | 0.94 (0.85, 1.03) | 0.97 (0.89, 1.07) | 0.95 (0.86, 1.04) | 0.50 |
| Multivariable+red meat | 1 (ref) | 0.95 (0.86, 1.04) | 0.94 (0.85, 1.03) | 0.98 (0.89, 1.07) | 0.95 (0.86, 1.05) | 0.57 |
| HPFS | | | | | | |
| Median, g/d | 4.3 | 5.4 | 6.3 | 7.4 | 9.2 | |
| No. of cases | 261 | 254 | 233 | 214 | 180 | |
| Age | 1 (ref) | 1.00 (0.84, 1.18) | 0.80 (0.67, 0.96) | 0.83 (0.69, 1.00) | 0.63 (0.52, 0.76) | <0.001 |
| Multivariable* | 1 (ref) | 1.01 (0.85, 1.20) | 0.82 (0.68, 0.98) | 0.86 (0.71, 1.04) | 0.67 (0.55, 0.82) | <0.001 |
| Multivariable+red meat | 1 (ref) | 1.01 (0.85, 1.21) | 0.83 (0.69, 1.00) | 0.89 (0.74, 1.07) | 0.73 (0.59, 0.89) | 0.001 |
| Insoluble fiber | | | | | | |
| NHS | | | | | | |
| Median, g/d | 9.5 | 11.8 | 13.7 | 15.8 | 19.6 | |
| No. of cases | 889 | 861 | 890 | 893 | 810 | |
| Age | 1 (ref) | 0.95 (0.87, 1.04) | 0.92 (0.84, 1.01) | 0.93 (0.85, 1.02) | 0.83 (0.75, 0.91) | <0.001 |
| Multivariable* | 1 (ref) | 0.95 (0.87, 1.05) | 0.93 (0.84, 1.02) | 0.94 (0.86, 1.04) | 0.86 (0.78, 0.95) | 0.004 |
| Multivariable+red meat | 1 (ref) | 0.95 (0.87, 1.05) | 0.93 (0.84, 1.02) | 0.94 (0.86, 1.04) | 0.86 (0.78, 0.95) | 0.005 |
| HPFS | | | | | | |
| Median, g/d | 10.7 | 13.5 | 15.8 | 18.5 | 23.1 | |
| No. of cases | 269 | 249 | 228 | 208 | 188 | |
| Age | 1 (ref) | 0.89 (0.75, 1.06) | 0.81 (0.68, 0.97) | 0.74 (0.61, 0.88) | 0.66 (0.55, 0.80) | <0.001 |
| Multivariable* | 1 (ref) | 0.90 (0.76, 1.07) | 0.82 (0.69, 0.99) | 0.76 (0.63, 0.92) | 0.71 (0.59, 0.86) | <0.001 |
| Multivariable+red meat | 1 (ref) | 0.91 (0.76, 1.08) | 0.84 (0.70, 1.00) | 0.79 (0.65, 0.95) | 0.77 (0.63, 0.94) | 0.005 |

* Soluble and insoluble fiber were estimated for 1990 and 1994 questionnaires in the NHS and 1986 and 1990 questionnaires in the HPFS.

* Further adjusted for body mass index, menopausal status and menopausal hormone use (women only), vigorous activity, alcohol intake, smoking, aspirin use, other nonsteroidal anti-inflammatory drug use, multivitamin use, acetaminophen use, physical examination, hypertension, hypercholesterolemia, calorie intake, and red meat intake.
